# Synthetic Data Generation and Random Projection Methods for Assessing Multivariate Similarity: A Low-Dimensional Methodological Study

**DOI:** 10.64898/2026.05.04.722226

**Authors:** John Heine, Erin Fowler, Steven Eschrich, Michael J. Schell

## Abstract

Data modeling in biomedical research often operates in the small-sample regime, where the number of observations is small relative to the data dimensionality; the detrimental effects of limited sample sizes are well documented in cancer studies. Synthetic data offers a potential solution to data shortfalls provided that the data generated is an adequate facsimile of the underlying distribution; the adequacy of such synthetic data remains an open-ended problem. In this work, we evaluate a synthetic generator proposed previously. We (1) develop a nonparametric method for assessing multivariate similarity based on the Cramér–Wold theorem and random projection testing, (2) investigate when the absence of bivariate correlation approximates independence in a non-normal setting, and (3) evaluate artifacts induced by data compression. The presentation is primarily methodological; low-dimensional data were used so each stage of the generation process could be analyzed explicitly. The generator applies a series of transformations to the observed data to accommodate the small-sample size, resulting in an uncoupled representation, where uncorrelated marginal distributions are modeled with optimized univariate kernel density estimation. Random projection outcomes were compared with a two-sample test and modeled as Bernoulli trials. Replicate outcomes were aggregated within each projection direction and pooled across directions, yielding a scalable standardized normal test statistic, under a working binomial approximation. The two-sample test significance level was decoupled from that governing the final normal inference. The projection framework also evaluated the full multivariate covariance structure. The generator produced high-fidelity multivariate synthetic data when bivariate correlation approximated independence in the non-normal setting. In highly compressed data, residual modes were best modeled as normally distributed regardless of their intrinsic distributional form. Ongoing work includes applying these methods to higher-dimensional, diverse data.

## 1. Introduction

Many biomedical datasets are characterized by a small-sample size (n) relative to their dimensionality (d), such that the observed data provides only sparse sampling of the underlying multivariate distribution. At the same time, advances in modern measurement technologies have made it possible to generate relatively large numbers of measurements per record, and the widespread use of modern analytical tools effectively enable unbounded exploratory modeling and hypothesis testing. The combination of small-sample size, increasing dimensionality, and effectively unrestricted exploratory analysis can undermine reliability, leading to well-known reproducibility challenges, including false discoveries [1] and overfitting [2]. To address these limitations (1) we develop a synthetic data generation framework tailored to the small-sample setting to support modeling and improved reproducibility, and (2) introduce a scalable, nonparametric method, for assessing multivariate distributional similarity, enabling rigorous evaluation of synthetic data fidelity.

Data modeling research requires sample sizes that can support both exploratory and reproducibility studies. In practice, however, these requirements are difficult to satisfy. Determining whether n is inadequate depends on the data dimensionality and its correlation structure. The type of modeling also plays a role in this determination. Typically, linear modeling requires smaller sample sizes than deep learning models. Nevertheless, small-sample size limitations are especially relevant to healthcare and biomedical research, where data availability can be restricted due to low-incidence diseases or underserved/underrepresented subpopulations [3], clinic visitation hesitancy [4, 5], inter-institutional data sharing obstacles [6], precision medicine costs [7], or study timeframes [8]. Detrimental effects from small-sample sizes are well documented in cancer studies [9, 10]. Studies with low statistical power not only have a reduced probability of detecting true effects but also tend to produce results that are less likely to reflect genuine relationships when statistical significance is achieved [11]. More broadly, the prevalence of non-reproducible preclinical research in life sciences has been estimated to exceed fifty percent, costing billions of dollars annually [12]. Accordingly, large-scale replication studies in preclinical cancer biology published in high-impact journals show that fewer than half of findings replicate and that estimated effect sizes are substantially inflated, consistent with instability in small-sample studies [13].

Synthetic data offers a potential solution to the small-sample size limitation. Recent work in synthesizing the available literature on the small data problem for machine learning concluded the future of big data may be in fact small data; these authors speculated for every thousand big datasets there are millions of small datasets [14]. This observation aligns with the discussion by Foraker, Mann, and Payne for the potential use of synthetic data in translational medicine [6]; relevant points from this article are summarized here. Treatment decisions optimized to the patient often require identifying patients with similar medical conditions and comparing outcomes across alternative therapies. Such analyses could be addressed by searching through medical records for patients with comparable conditions. However, archives of this kind are usually rarely available in the routine practice setting. Although data sharing could help mitigate this problem, practical mechanisms for doing so are limited. As reviewed by these investigators [15], machine learning can support precision medicine by detecting complex patterns within the data enabling patient-level prediction; however, progress is limited by small-sample sizes, heterogeneous datasets, confounding variables, and reproducibility challenges. The authors have worked in biomedical-healthcare cancer research for many years within a Comprehensive National Cancer Institute-designated Cancer Center and have encountered this persistent challenge of insufficient sample sizes over decades.

Synthetic data offers a potential solution to small-sample size limitations provided the data is an adequate facsimile of the underlying distribution given a representative but small-sample size. However, defining what constitutes adequate facsimile remains an open-ended question. These researchers [16] describe simulation studies where datasets are generated from specified probability distributions or empirical data-generating mechanisms to evaluate statistical methods; however, these generators assume the underlying distribution is known and therefore do not address the more general problem of generating realistic data when the true multivariate distribution is unknown. Existing synthetic data generation applications in healthcare often focus on generating records from large populations [17–20]. For example, generalized adversarial network (GAN) based methods have demonstrated impressive capabilities, particularly in image generation tasks, and have increasingly been applied to tabular data and other domains [21]. However, notwithstanding their success, evaluating whether synthetic data generated by GANs or other generative models faithfully reproduces the statistical structure of the original dataset remains a challenging problem; existing evaluation strategies often rely on indirect measures such as predictive performance or summary statistics, which may not fully capture differences between multivariate distributions [21]. A recent review accompanied with extensive analyses [22] evaluated four general purpose synthetic data approaches [23–26], considered the most influential; the evaluation found considerable variation in performance. These authors [22] concluded more work is required in evaluating such generators, a finding echoed by other recent reviews [21, 27]. A commentary and survey of synthetic data generation approaches also concluded that the inadequate sample size problem has received little attention [28], paralleling similar observations [29].

Direct comparison of multivariate distributions is challenging. Many classical goodness-of-fit approaches require reliable estimates of the underlying probability densities. In practice, this often involves constructing multidimensional histograms or kernel density estimates. However, when sample sizes are limited, even in relatively low dimensions, the data occupy only a sparse portion of the multivariate space as illustrated explicitly, previously [30]. As a result, multidimensional histograms contain large numbers of empty or poorly populated bins, making stable density estimations and subsequent divergence calculations unreliable or impractical [31]. Consequently, direct density-based comparisons become difficult to apply in small-sample size settings. Thus, the non-parametric random projection test is developed into a formal statistical inference framework.

Our goals of generating synthetic data and developing methods for assessing its validity are under development specifically to address the non-normal, small-sample-size problem [30]. The generation of realistic synthetic data can address important challenges in applied biomedical engineering and is also an interesting data-analytic problem on its own. Two closely related problems remain in synthetic data generation: (1) can an approach produce high-fidelity data when only a small but representative input dataset is available, and (2) what metrics are most appropriate for evaluating the validity of such synthetic data? Our approach addresses both problems: synthesizing the population distribution from a small representative sample and evaluating the resulting synthetic data. The proposed method is modular, with specific assumptions underlying each stage of the generator. In tandem, we are developing scalable methods to evaluate the authenticity of the synthesized data.

In this report, we investigate when the absence of bivariate correlation approximates independence in the non-normal setting and develop methods for assessing multivariate similarity using random projection testing. We also investigate a counterexample, where the data are compressed, with dominant marginal modes well represented as normal; the residual (i.e., a component in the uncoupled space that explains an inconsequential fraction of the variance) was investigated in more detail than previously [30]. In this current work, low-dimensional data were intentionally selected to allow intermediate analyses and dependence structures to be examined directly. The observed data is transformed into two additional representations, enabling explicit comparison of the data and their dependence structures across all three representations. The purpose of this work is therefore methodological: to demonstrate the behavior of the proposed synthetic data generator and to refine a projection-based comparison test in a controlled setting before extending the framework to higher-dimensional problems with limited observations. In our previous work, we demonstrated that observed data could be reconstructed using this approach, provided the data exhibit latent normal behavior in higher-dimensional settings [30]. Building on this foundation, the methods developed here introduce an additional level of complexity to both the synthetic data generator and the multivariate comparison framework. This work gives a strong emphasis on advancing the random projection analysis into a standardized statistical testing framework and examining its behavior in greater depth. We show that this nonparametric test not only assesses distributional agreement, but also implicitly evaluates the underlying multivariate covariance structure. We also investigate the test behavior under various conditions to better understand its output metric.

## 2. Methods

### 2.1 Synthetic data generator

We use the following conventions and notations in this report. The normal form is used to compute covariance matrix elements. A normalized empirical histogram is referred to as an empirical probability density function (pdf) or distribution. E(·) is the expectation operator, and var(·) denotes the variance of a given quantity. Components of a vector (dimensions of size d) are referred to with an integer index or subscript ranging from 1-d; all vectors are column vectors with d-components unless specified otherwise and designated by lower-case bold fonts (e.g., **x**). An uppercase bold font denotes a matrix (e.g., **V**). The term comparison is reserved for evaluating one dataset against another dataset or collection of datasets. Each dataset is comprised of n observed or synthesized records, where each record is represented by a specified d-dimensional vector.

A brief overview of the generator is provided first following previous work [30]. The observed dataset initially exists in the X representation, where n observations (records) are represented by d-dimensional vector **x** with covariance matrix **C**_x_. The analysis proceeds through multiple steps by involving transformations of observed **x**; each component of **x** is transformed to a standardized zero-mean unit variance normal variate producing the Y representation and vector **y** with covariance **C**_y._ Next, an orthogonal PCA transformation is applied to **y**, producing the T representation and vector **t** with diagonal covariance **C**_t_. These steps are explained explicitly below. A dataset consists of n observed or synthesized records each represented by a corresponding vector (e.g., **x**, **y**, or **t**).

**Step 0**. Observed **x** is first mean centered (still labeled **x**) and compared with multivariate normality: (1) univariate marginal pdfs are compared with normality; and (2) if all marginal pdfs satisfy normality, multivariate normality is then evaluated. If **x** satisfies multivariate normality, standard multivariate normal generation techniques can be applied (i.e., the problem is essentially solved). Often normality can be rejected by simple observation when the dimensionality permits, as measured variables often have right-skewed marginal pdfs (such measurements often have zero probability of being negative). When observation fails or is not practical, the Kolmogorov-Smirnov (KS) test is applied using the normal distribution as the comparator with a 0.05 significance level for marginal testing. However, if **x** fails to obey multivariate normality, the algorithm proceeds to the next step.

**Step 1**. This stage specifically addresses the small-sample size problem. Each component of observed **x** is treated independently and modeled using optimized univariate kernel density estimation (KDE) with a normal kernel. Bandwidth selection for each component is determined through d independent optimization procedures using differential evolution [32]. In principle, the resulting distributions allow the sample size n to be expanded arbitrarily. Synthetic records (realizations) generated in this step are used only to augment the observed data to construct smooth, nearly continuous univariate transformations for the X-Y transition. Here, observed and synthetic pdfs are compared with the KS test as the fitness metric in the differential evolution optimization procedure described previously [30, 33].

**Step 2**. Each component of **x** (observed only) is transformed into a standardized normal variate with zero-mean and unit variance. This transformation produces the Y representation, where each observation is described by the vector **y** with covariance matrix **C**_y._ Synthetic data generated in the X representation are used only for constructing the mapping functions to ensure smoothness, near continuity, and improved estimates of the dynamic range for each component. These synthetic realizations are discarded after the mapping is performed because they do not preserve the canonical covariance structure of the original data. There is no guarantee that **y** is multivariate normal; hence, **y** is tested for multivariate normality. When **y** is multivariate normal, the original data in X can be interpreted as arising from a latent multivariate normal representation. In any event, the analysis proceeds to the next step.

**Step 3**. The observed dataset in Y is projected into the T representation using principal component analysis (PCA); the PCA transformation is determined with the observed **y**. This transformation produces the vector **t** with diagonal covariance matrix **C**_t_. When **y** is multivariate normal, the marginals in T will also be normal because a linear transformation of a multivariate normal process produces another multivariate normal process. In this case, synthetic data can be generated by sampling independent zero-mean normal random variables for each component of **t**, where the variances are the respective eigenvalues with virtually unlimited realizations. The inverse PCA transformation then restores the canonical covariance structure in Y, and the inverse Y to X transformations restore the original coordinate system. The same process is used to generate a normal multivariate comparator in Y; thus, for a given dataset, a multivariate normal comparator is obtained, noting the inverse PCA transformation restores the canonical covariance, **C**_y_. Then, applying the inverse Y to X transformation to each component of **y** provides a latent multivariate normal comparator dataset characterized in the X representation with vector **x**.

### 2.2 Objectives

In our previous work, we showed that the T representation can also be used to perform diagnostics on **y** by testing the univariate marginals of **t** for normality. Multivariate normal synthetic data in T was generated by approximating the marginals of **t** as independent normal variates using the Box–Muller method [34]. We found this approach worked well when marginal distributions in T that deviated from normality explained only a small fraction of the total variance (≪1%); in these situations, the residuals were modeled as normal, even though statistical testing showed significant deviations from normality. In this prior work, we did not encounter situations where the dominant modes in T deviated significantly from normality [30].

The work presented in this report is primarily methodological and comprises three related advancements (i.e., objectives): (1) *investigate when the absence of bivariate correlation approximates independence in a non-normal setting*. Here, we investigate the case where a non-negligible marginal distribution in T is not well approximated as normal and determine if the multivariate distribution can still be synthesized by modeling the univariate marginals with KDE or KDE in combination with normal marginal models. Because the covariance matrix in T is diagonal, this coordinate system provides a convenient framework for constructing multivariate synthetic data from independently generated marginal distributions; (2) *evaluate artifacts induced by data compression*. Here, we study the case where dominant modes are normal in T and the residual is non-normal. This case replicates similar situations encountered in prior work [30]. Here, we investigate the residual in more detail to understand the appropriate modeling approach; and (3) *develop a nonparametric method for assessing multivariate similarity and random projection testing*. For this advancement, we extend a non-parametric random projection test for multivariate normality developed previously to the more general framework of comparing two multivariate distributions in support of the generation process. We derive a standardized statistical test starting with the Cramér-Wold theorem [35] for multivariate similarity. The resulting method provides a new two-level testing architecture that converts repeated nonparametric distributional comparisons into a tractable standardized normal test statistic, under a working binomial approximation. We also show how the same random projection test also interrogates the covariance structure under various conditions. The projection test development is provided with full technical detail to support independent review, validation, and replication. Developing this test supports the analysis in the generator steps outlined above. Low-dimensional datasets were intentionally selected so that the intermediate analyses could be examined directly. This allows the transformations between the X, Y and T representations and data dependence structure to be observed explicitly.

### 2.3 Study Data

Data used in this report was derived from two sources: bean morphology and mammographic images, each analyzed with d = 3 variables. The low dimensionality was selected deliberately to isolate the study so that each step of the proposed procedure could be directly observed and evaluated. The study was not conducted in the small-sample regime, thereby reducing the possibility that the observed behavior was an artifact of small-sample effects. Although data from these sources were used in our previous work, the specific variables analyzed here differ from those used previously.

Bean data is open source and comprised of seven bean types [36]. A bean type was selected randomly from those with at least n = 1334 observations (5 of the 7 types satisfied this condition), resulting in selection of the Horoz bean class with n = 1978; from this, a subset of n =1334 bean-observations were selected randomly without replacement. The full population contains 16 measurements per bean. From these d = 3 variables with at least moderate linear correlation were selected: x_1_ = perimeter, x_2_ = aspect ratio, and x_3_ = roundness (defining the observed bean dataset). The number of observed records was restricted to match that of the mammography dataset for cross dataset comparison purposes.

Mammography data consists of measurements derived from mammographic images (n = 1334). Three quantities were measured from within the breast region: x_1_ = breast area (cm^2^), x_2_ = standard deviation of the pixel values, and x_3_ = standardized skewness of the pixel values, related to the third central moment (defining the observed mammography dataset). These images and other non-image data are also opensource [37]. Each image was from a different study participant.

All procedures were carried out in accordance with the relevant guidelines and regulations approved by the Institutional Review Board of the University of South Florida, Tampa, FL under protocol 104715_CR000002. Data was collected both prospectively and retrospectively over these dates: 24/10/2006 to 21/10/2019. Prospective data was collected with written informed consent. Retrospective data was collected with a waiver of informed consent provided by said IRB. The data was last accessed for research purposes on 01/02/2026. No minors were included in the study. Two authors (JH and EF) have access to identifiable individual participant data.

Synthetic data refers to vectors (e.g., three-component vectors) randomly sampled from the corresponding synthetic population, which is assumed to be of effectively unlimited size. For comparisons with normality or with the observed datasets, a synthetic dataset denotes a collection of n = 1334 vector realizations (records) drawn from this synthetic population (i.e., synthetic **x**, **y**, or **t**). For comparisons in X, Y, and T, the representative vectors of a given dataset were transformed accordingly.

### 2.4 Univariate Comparisons

First, the observed univariate pdfs were compared against normality. Second, in each representation, synthetic and observed univariate marginal pdfs were also compared. In both cases, one thousand synthetic **t**, **y**, or **x** datasets were generated (each with n = 1334) and compared with the observed dataset. The KS test was used for all comparisons with two-sided significance level α = 0.05. The percentage of times that the null hypothesis was not rejected was recorded.

### 2.5 Multivariate Random Projection Test

To avoid these limitations discussed above, we are developing a random projection approach that reduces the multivariate comparison problem to scalar (distributional) comparisons. The approach is motivated by the Cramér-Wold theorem [35] that states two multivariate distributions are identical if and only if all one-dimensional projections of the distributions are identical. The test is built on generating random projection vectors. The inner product of a given random projection vector (i.e., a vector that points in a d-dimensional random direction) with a data vector (observed or synthetic) produces a scalar quantity; the distributions of these scalar quantities (observed versus synthetic datasets) were compared with a two-sample test (i.e., the KS test). Many synthetic datasets are compared with the observed data for a fixed direction and then over many random directions. Through repeated projection comparison outcomes, a standardized test is developed. We also show that the same framework tests for covariance similarities. The development starts in the T representation because the covariance matrix is diagonal, which simplifies the analysis of dependence among variables, but the test applies equally as well in Y and X as demonstrated. Thus, multiple synthetic datasets are generated in T and compared with the respective observed dataset. The analysis is also performed in Y and X when reversing the steps.

The random projection test is formally developed in this section. Let **t** ∊ R*^d^* define the vector of principal component variables for a given dataset. A given d-dimensional random projection vector **a** is generated with components randomly sampled from a standard normal distribution and then normalized to unit norm

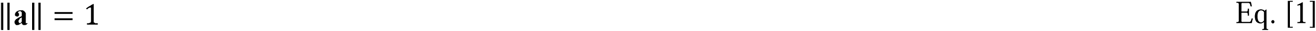

For a given projection vector **a**_k_, the scalar variable from the projection is computed as an inner product:

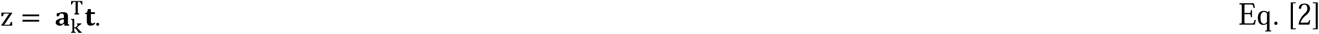

When two datasets are compared, projected records from each dataset are obtained using the same projection vector. The resulting distributions from the projections are then compared using the KS test (i.e., the two-sample test), which evaluates the null hypothesis that n projected records obtained from each dataset are drawn from the same distribution. Here, the two-sided significance level α = 0.05 was used. In our previous work, heuristics were employed to derive the projection test fidelity, and the test was developed for normality comparisons [30]. Here, these operations are constructed into a formal nonparametric testing framework for comparing two multivariate distributions. Derivations are used to show how this test also interrogates the covariance structure, which are provided after the test-statistic development.

Because a single projection captures only limited information about the multivariate structure, the procedure is repeated over many independent projection vectors to approximate the Cramér-Wold condition. The test is built from the inside out by first comparing the observed dataset with many synthetic datasets for a fixed direction (replicates) and then by aggregating the outcomes across many random projection directions. Let K define the number of random projection directions and R the number of replicate comparisons for each direction giving N = K × R tests (i.e., R synthetic dataset realizations are compared with the observed dataset for a fixed direction k and then over K directions). For each **a**_k_ projection, a KS test is performed with the observed **t** against multivariate synthetic **t** realizations. For each replicate dataset r, a corresponding synthetic vector denoted by **w**_i,r_ is compared with the observed **t** via projections. For the projection direction **a**_k_, this operation is performed for the observed dataset giving

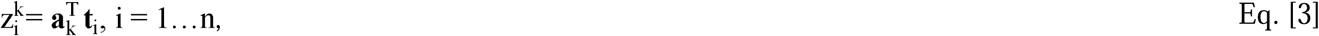

where the index i has been introduced as the record vector index (n =1334). For the same projection direction, the synthetic vectors in T, relabeled as **w** to avoid confusion, are projected similarly.

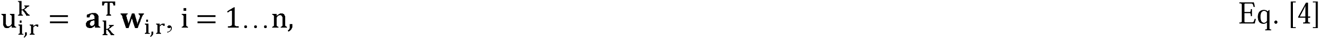

where the index r is included to indicate multiple replicate synthetic datasets for comparison with fixed k; thus, there are R synthetic dataset comparisons with the observed for each of the K random directions.

Next, we use the above KS test comparisons to develop a statistical testing framework. The technique is motivated by considering the series of N = KR comparison outcomes as a sequence of binary outcomes representing successes and failures. For each projection direction k, the projected observed and synthetic dataset distributions are compared, producing R replicate outcomes per direction. Let 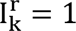 when the null hypothesis is not rejected for each replicate r comparison in direction k, or 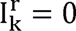 otherwise. Under a working binomial approximation, we model the R replicate outcomes for a fixed direction as independent Bernoulli trials with probability of success p so that the number of successes in each direction follows a binomial distribution. In this context, the KS test is used as a local discriminator for each projection, with its nominal significance level (α = 0.05). Extending across all projection directions (i.e. summing K binomial variables with a common success rate), the total number of successes

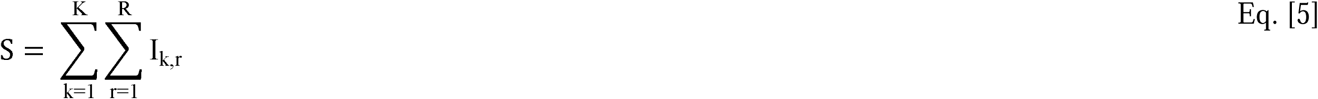

is modeled as a binomial random variable with parameters N = KR, success probability p, and failure probability 1-p with

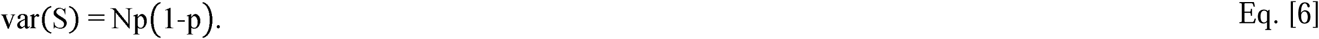

We define the random test-statistic Q as the related proportion:

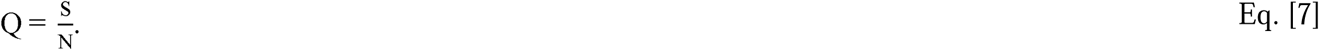

Using this random variable transformation gives

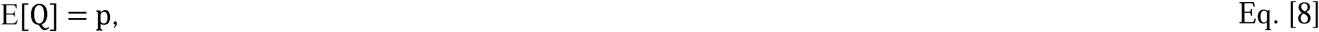

and

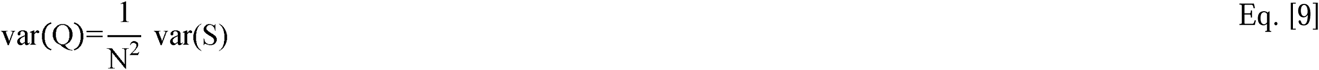

yielding

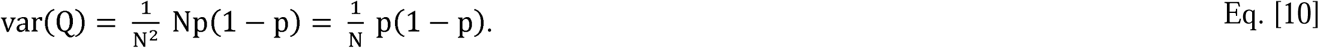

Taking the square root of Eq. [10] gives the standard deviation (SD) for Q:

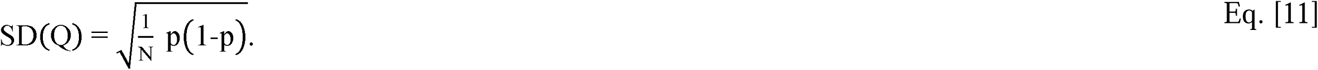

For large N, the binomial distribution is well approximated as normal:

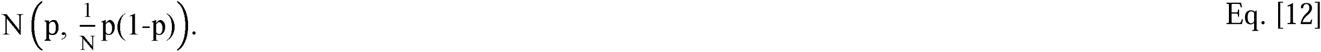

Thus, the standardized test for Q (i.e., the Q-test) is specified as

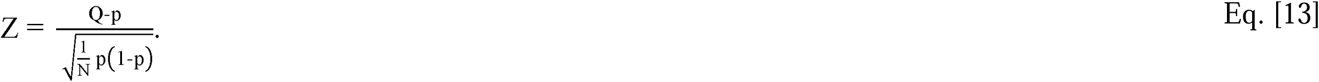

Consequently, the test-statistic 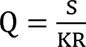 is the fraction of simulations where the observed and synthetic dataset distributions are in agreement (over all directional replicates and directions) relative to the design factor p. The critical value for Z is derived from a standardized normal distribution at a specified significance level that is distinct from the per-projection α level from the KS test comparisons. The hypothesis test for Q is thereby specified as

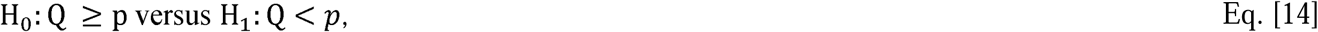

where p is the probability that a projection comparison does not reject the KS null hypothesis under the working model, and Q is the observed quantity. In practice, we prespecify p_0_ as a target for the fraction of projection comparisons where the KS null hypothesis is not rejected (here, p_0_ = 0.90). We set p = p_0_ in Eqs. [13 and 14] and compare the observed Q with a nominal lower-tail cutoff derived from the standard normal approximation at a one-sided level of 0.05. This level is distinct from the significance level of each KS comparison. Because dependence may exist among projection outcomes, this level is conditional on the working binomial approximation and is not an empirically calibrated error rate. Likewise, the same test applies in Y, and X. Moreover, the analogous Q-test would result when an arbitrary experiment comprised of M comparisons are modeled as independent Bernoulli trials with success rate p and failure rate 1-p. Following the development leading to Eq. [13], p is the mean, and p(1-p)/M is the variance. These two quantities produce the analogous Q-test.

The use of multiple projection directions and synthetic replicates serves two purposes. First, using R synthetic dataset replicates reduces variability associated with the stochastic synthetic process and provides a stable estimate of the projection agreement probability for a fixed direction. Second, using many randomly sampled projection directions, K, provides a stochastic approximation to the Cramér–Wold characterization of multivariate distributions. Under the working binomial approximation, the precision of the estimator Q increases with the total number of projection comparisons N = KR, with variance proportional to 1/(KR). Consequently, small deviations between the observed and synthetic projected distributions become detectable for large N. In this study, we use K = 1000 directions with R = 1000 replicates, yielding 10^6^ projection dataset comparisons, ensuring a stable estimate of distributional agreement.

The primary analysis is performed in the T representation for these reasons. Principal component analysis diagonalizes the covariance matrix, producing orthogonal variables with variances ordered by their contribution to the total variance. This representation simplifies the data dependence structure and provides a convenient framework for investigating whether the absence of correlation implies independence. Even when the marginal pdfs in T are approximately normal, this does not guarantee the joint distribution is multivariate normal. Because uncorrelated variables are independent only for the multivariate normal distribution, the independence assumption in T must be verified empirically. For non-normal data, higher-order dependence may remain. Therefore, independent marginal pdf modeling in T using univariate KDE is not guaranteed to reproduce the full multivariate distribution. The random projection comparison test is therefore used as a diagnostic to determine whether synthetic data generated under this assumption adequately reproduces the observed multivariate distribution.

When the projected comparisons indicate multivariate agreement in T, the results are subsequently verified in the Y and X representations to ensure that the conclusions remain consistent in the two other correlated coordinate systems.

### 2.6 Random Projection Test and Covariance Structure Interrogation

Next, we show that the same random projection test also probes the covariance structure of the interrogated distribution starting in Y; as above (z is a generic variable) for a random direction (with unit norm) the projected scalar is

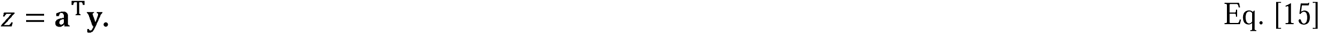

The expected variance of the projection z (distributional sense) in Eq. [15] is expressed as

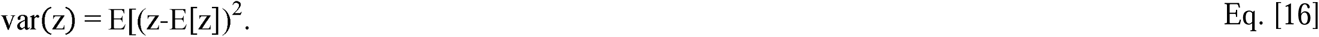

Equivalently,

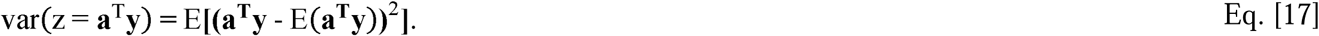

Because the expectation operator is linear, working from the inside of the quadratic in Eq. [17] gives

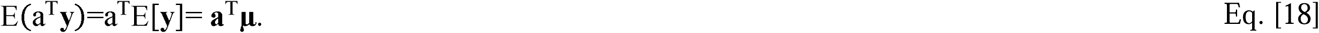

Although, in this work **µ** is identically zero in Y. Factoring the quadradic term in Eq. [17] gives

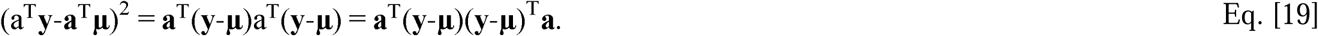

Recognizing the central factor as the covariance structure for **y** and using the linear property of the expectation gives

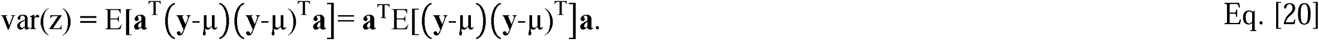

Taking the expectation in Eq. [20] yields

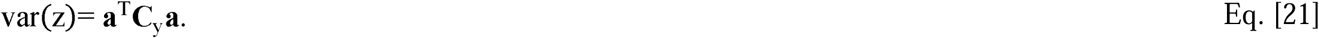

To show how this relates to the analysis in T, the PCA decomposition defines

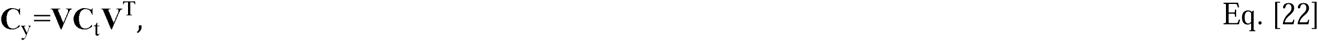

where the columns of **V** are the normalized eigenvectors of **c**_y_. Substituting Eq. [22] into Eq. [21] gives

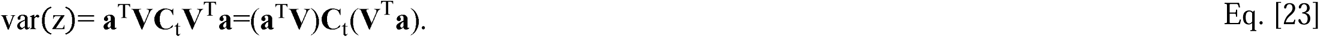

Making the substitution

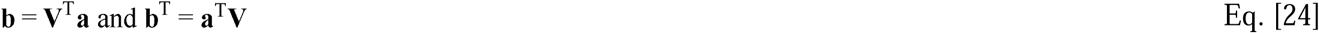

in Eq. [23] yields

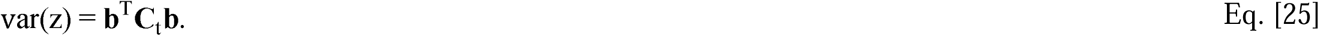

Because the covariance is diagonal, this gives an expansion with the components of **b**

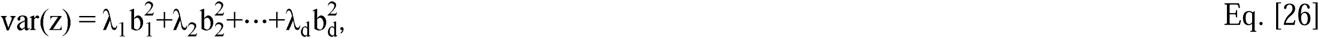

where λ_i_ are the eigenvalues of **C_y._** Equation [26] can be expressed equivalently noting that

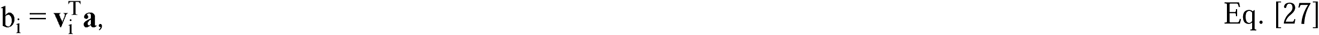

where **v**_i_ (columns of **V**) is the i^th^ eigenvector in **V**, yielding

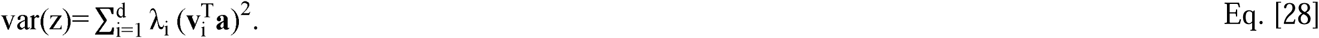

The variance of a random projection **a** is therefore a weighted mixture of PCA eigenvalues with these weights

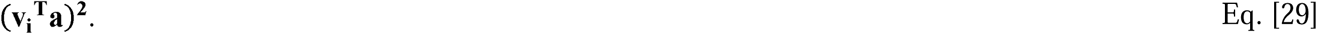

Each weight measures the strength of how the projection direction aligns with each PCA axis. Because both the eigenvectors and **a** have unit norm, Eq. [28] is expressed equivalently using the inner product identity

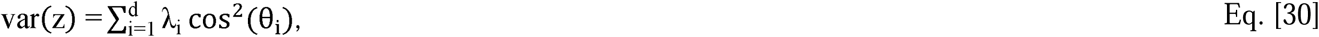

where θ_i_ is the angle between **a** and **v**_i_. Therefore, random projections sample mixtures of eigenvalues and implicitly test the geometry in T. Thus, the square of the inner product term in Eq. [30] is the fraction of the projection variance coming from the i^th^ principal component. We note that the test originated from a random projection in Y, where projected variance reflects the underlying covariance structure (see Eq. [23]) and has an equivalent interpretation in Eq. [30] as probing the data in T through mixtures of eigenvalues. Consequently, the sampled random projections probe aspects of the multivariate distribution, including its covariance structure, without explicitly estimating a multivariate density.

### 2.7 Random Projection test and Coordinate System Equivalence

The connection between the projection test in Y and T is shown more explicitly by starting in T with the normalized projection vector **g** giving (z is generic and not equivalent with z above directly)

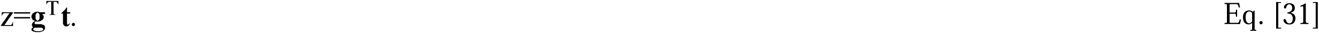

Because

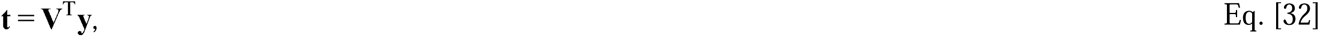

then

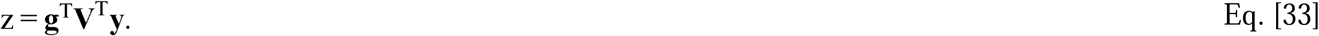

Expressing Eq. [33] equivalently shows

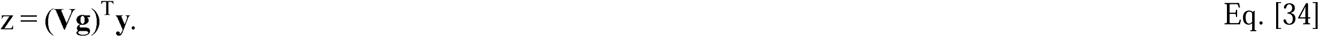

Thus, applying projection vector **g** in T is equivalent to applying vector **a** = **Vg** in Y; this is the same projection vector rotated into the other coordinate system or vice versa. Equation [34] explicitly illustrates the equivalence of the projection test applied in either T or Y. In general, the equivalence does not carry over to X because the transformations from Y to X may be non-linear and are performed univariately. In this situation, the test must be applied empirically to show distribution equivalence in X.

### 2.8 Random Projection Test and Scaling

Special attention is required when using the projection test in X or in situations where vector components are not mean centered. Here, we point out two more obvious conditions. Observed data in X is not mean centered in our case (as observed) and in many situations. When evaluating non-centered data, the projection test is valid but may produce results with the incorrect interpretation. First, if data in X is positive valued for univariate marginals, **x** lives in a restricted quadrant in d-dimensional space, whereas the projection vectors span the entire space, giving one reason for misinterpretation. Second, there is another condition with at least equivalent gravity. The projection test in X (or any setting where this principle applies) is expressed as

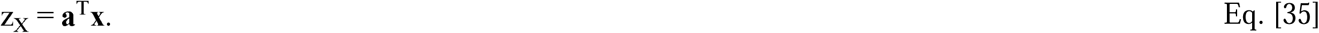

Expressing **x** equivalently gives

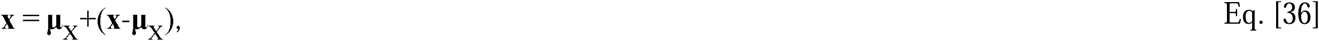

where

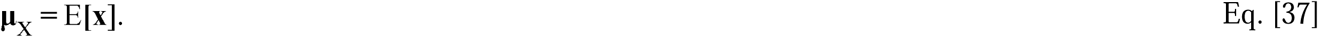

The equivalent projection for vector **a** is

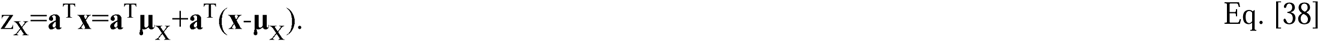

On the right-hand side of the equal sign in Eq. [38], the projection of the mean vector **μ**_X_ produces offset contributions and second projection term describes random variation. In the case where a given component of **μ**_X_ is large relative to the respective variation, a random projection may become biased in given direction. Both conditions imply mean centering should be applied before employing the projection test. Evaluating projections in Y might be the most appropriate setting to compare multivariate similarity; the marginals have equal representation in the mean, variance, and other moment characteristics, although the data was measured and lives in non-centered X representation. Although scaling is recommended, conversely a variable with the largest value may be the more important variable, or one with the least measurement imprecision; in such a situation, biasing a test in that direction may be appropriate, if interpreted correctly.

## 3. Results

### 3.1 Univariate analysis

Univariate descriptions of observed **x** (bean and mammography datasets) are provided in Table 1a; in X and Y, these were derived from the multivariate reconstruction starting in T (Model C for the synthetic bean datasets, described below). Also included are the PCA explained variances in T and marginal pdf comparisons with normality for the observed datasets. In X (original space), the bean dataset marginal pdf for x_3_ deviated from normality, and all marginal pdfs deviated significantly from normality in the mammography dataset (see p-values). As expected, in Y (standardized univariate pdfs) all marginals were not significantly different from normal (see p-values). For **t** (PCA-transformed), the bean dataset is essentially compressed into two dimensions. The dominant two modes explained approximately100 percent of the variance and were well approximated as normal, whereas the residual component showed significant deviation from normality. For **t**, the mammography dataset exhibited relatively less compression in comparison with each mode explaining a considerable percentage of the variance; the first and third modes explained about 45 and 22 percent of the variance, respectively, and showed close agreement with normality, whereas the second mode explained about 33 percent of the variance and deviated significantly from normality. Figure 1 shows observed bean dataset marginal pdfs (blue): T (top), Y (middle), and X (bottom). The mammography dataset pdfs are shown in Figure 2 in the same arraignment.

**Table 1a.**
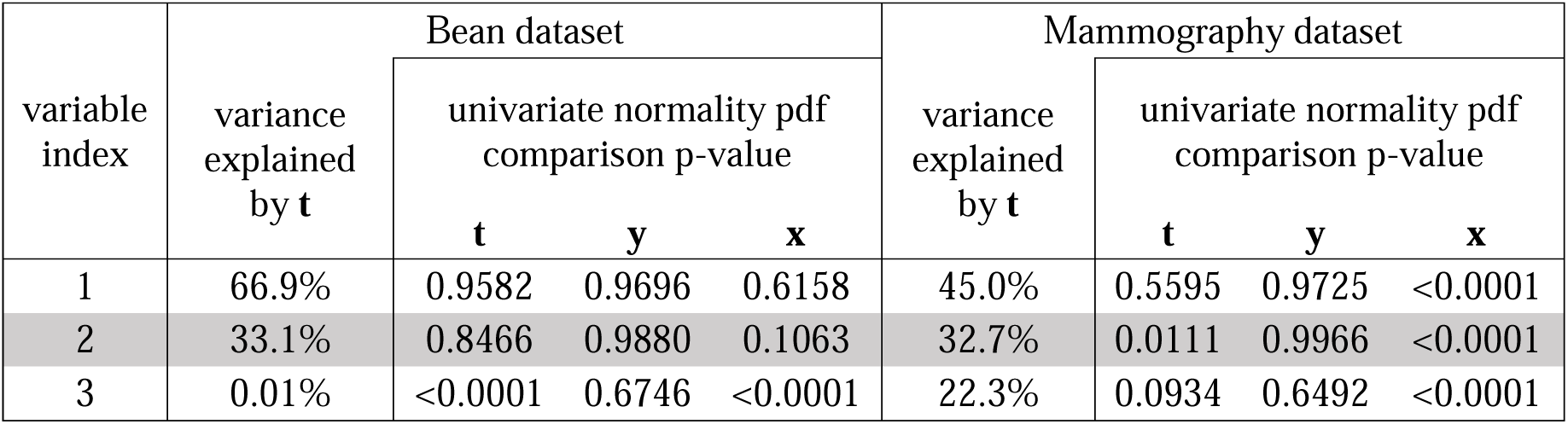
Univariate marginal pdf normality comparisons between the observed bean dataset (left) and mammography dataset (right): the percentage of the variance explained for each component of the vector **t** is provided for each dataset. Each univariate marginal pdf was compared with normality. The p-values from the KS test are provided in the respective **x**, **y** and **t** columns.

**Figure 1.**
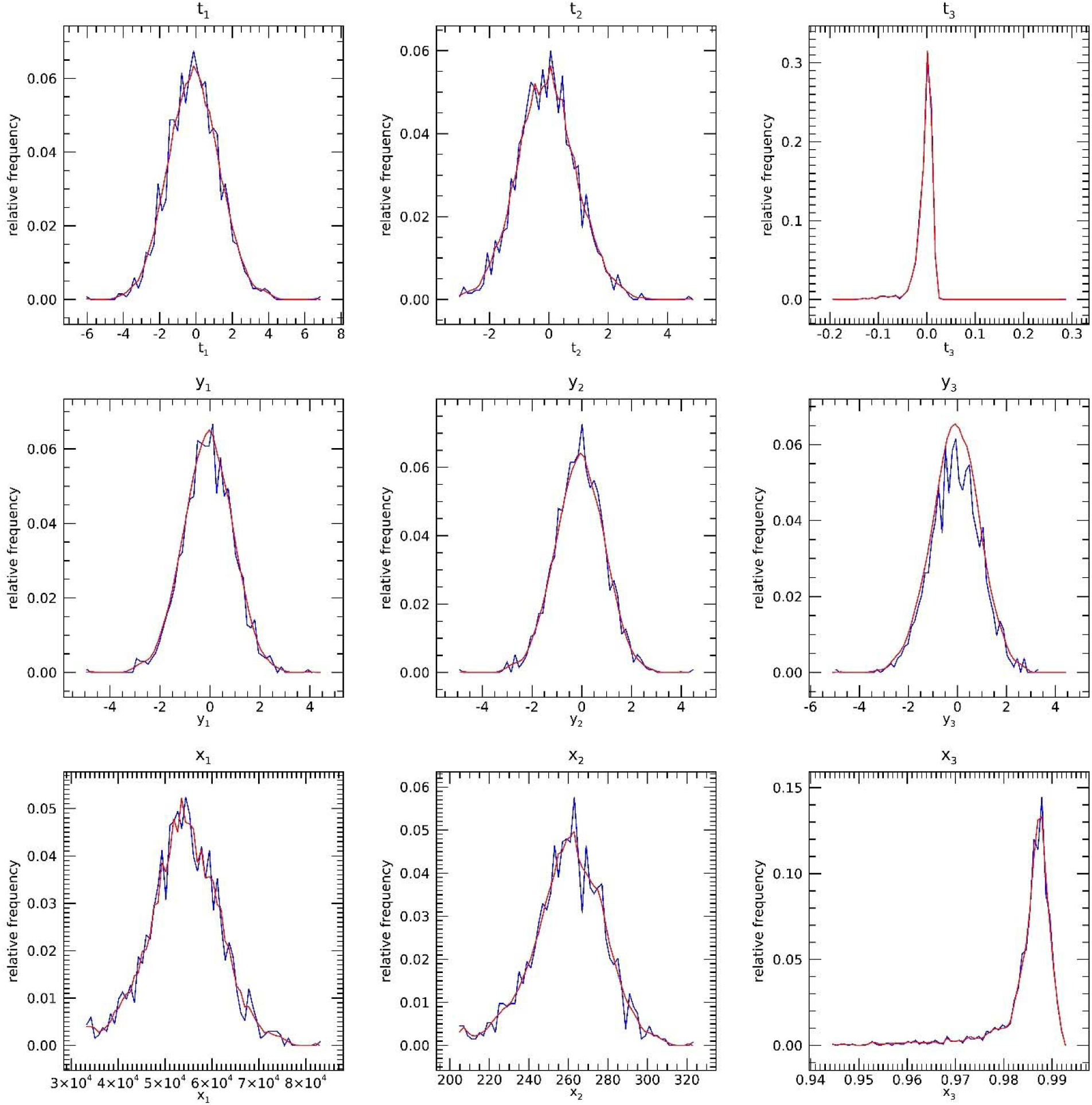
Observed and synthetic bean dataset marginal pdfs: observed dataset pdfs (blue) are compared with synthetic dataset pdfs (red) derived from a random realization: **t** (top), **y** (middle), and **x** (bottom). The index indicates vector components. Synthetic data was generated with Model C starting in the T representation.

**Figure 2.**
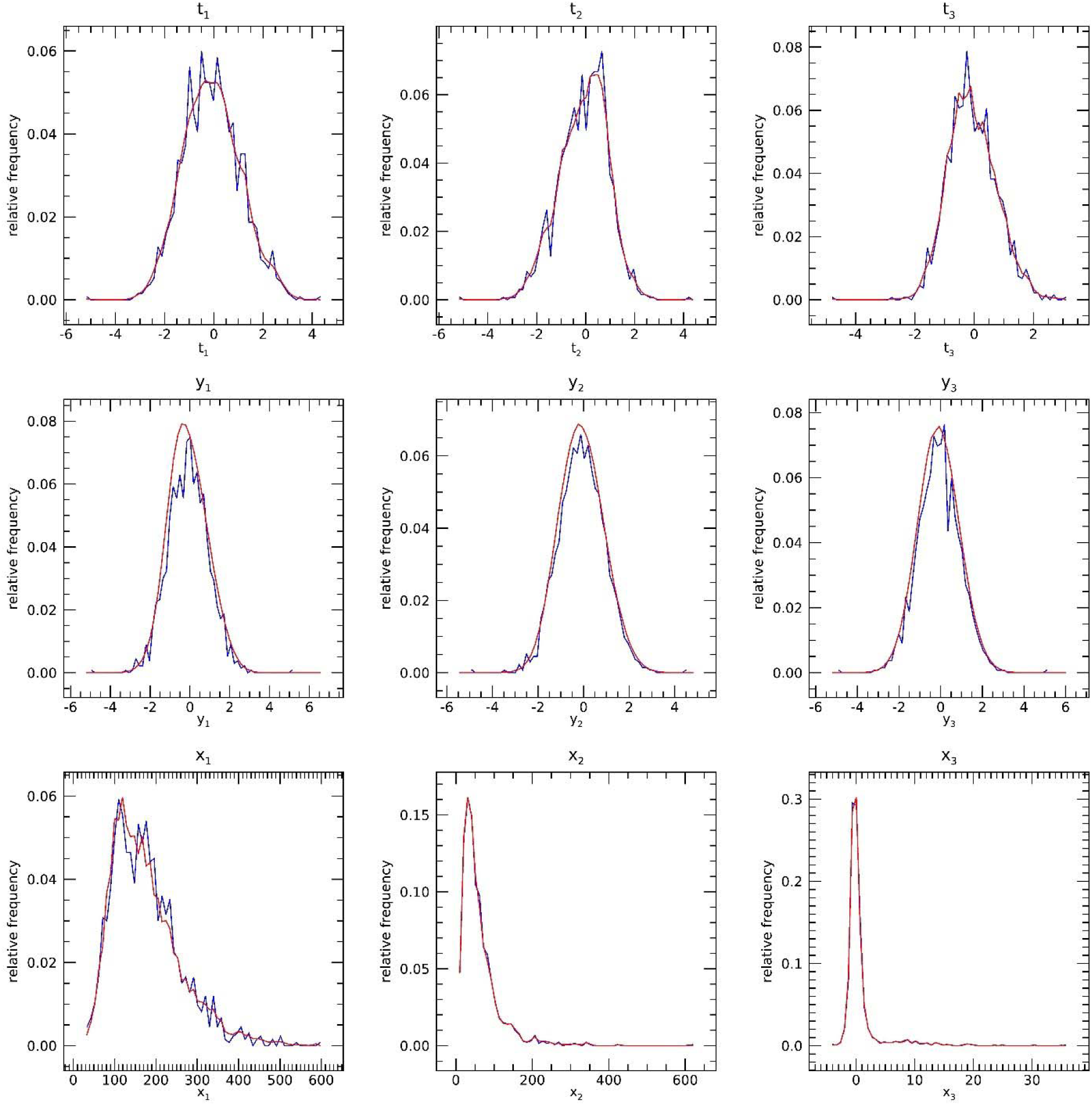
Observed and synthetic mammography dataset marginal pdfs: observed dataset pdfs (blue) are compared with synthetic dataset pdfs (red) derived from a random realization: **t** (top), **y** (middle), and **x** (bottom). The index indicates vector components. Synthetic data was generated with univariate kernel density estimation starting in the T representation.

Next, comparisons between univariate marginal pdfs are investigated between the observed and synthetic datasets. Figure 1 (bean dataset) and Figure 2 (mammography dataset) show comparisons of marginal pdfs between the observed (blue) and synthetic datasets (red): (1) T (top, (2) mapped from T to Y (middle), and (3) inverse mapped from Y to X (bottom). Here, one synthetic dataset (shown in three representations) was selected at random for each observed comparison. For both the bean and mammography datasets, the observed and synthetic marginal pdfs show close agreement visually. Table 1b details the univariate observed and synthetic pdf comparison results as the fraction of times the null hypothesis was not rejected by the KS test. Most comparisons were greater than 0.95, showing strong univariate agreement. As a heuristic criterion, we use less than 0.90 as the cutoff for agreement. These proportions can be used as test-statistics in a more formal Q-test inference, discussed below.

**Table 1b.**
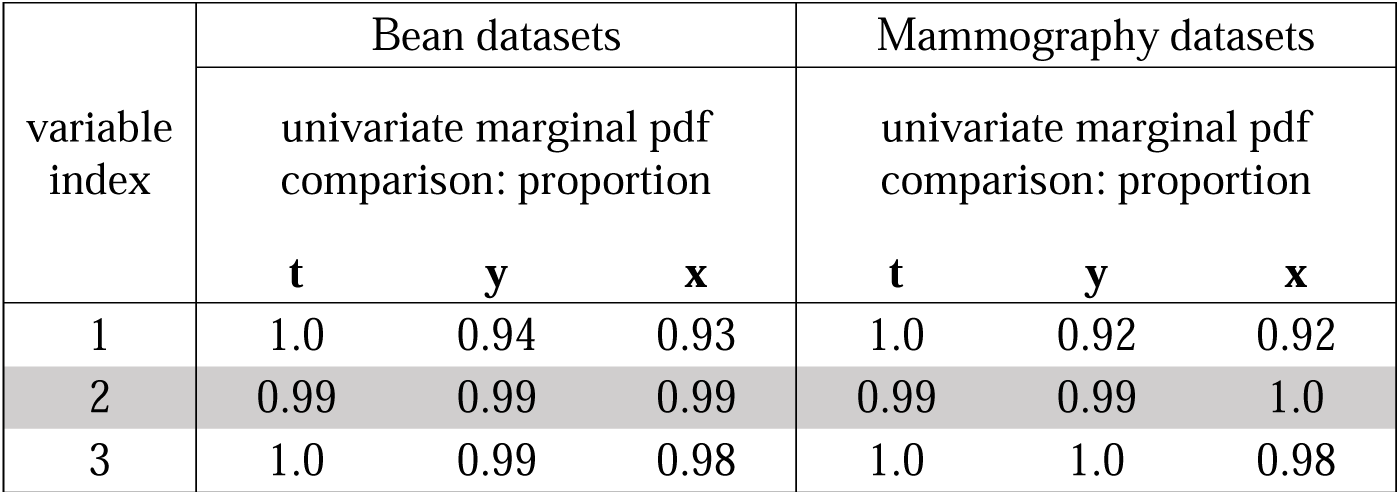
Univariate marginal pdf comparisons between observed and synthetic datasets: each univariate marginal pdf from the observed dataset was compared with 1000 synthetic dataset realizations. The proportion of times the null hypothesis was not rejected from the KS test values are provided under the **x**, **y** and **t** columns. The critical threshold Q_c_ ≈ 0.88,

| variable<br>index | Bean datasets |  |  | Mammography datasets |  |  |
| --- | --- | --- | --- | --- | --- | --- |
|  | univariate marginal pdf<br>comparison: proportion |  |  | univariate marginal pdf<br>comparison: proportion |  |  |
|  | **t** | **y** | **x** | **t** | **y** | **x** |
| 1 | 1.0 | 0.94 | 0.93 | 1.0 | 0.92 | 0.92 |
| 2 | 0.99 | 0.99 | 0.99 | 0.99 | 0.99 | 1.0 |
| 3 | 1.0 | 0.99 | 0.98 | 1.0 | 1.0 | 0.98 |

### 3.2 Q-test and Multivariate analysis

The analytical multivariate similarity with normality and comparisons with the observed and synthetic datasets were investigated next. For the Q-test (addressing the empirical component of objective 3) we set the agreement parameter p_0_ = 0.90. Calculating the standard deviation for Q with N and p_0_ (using Eqs. [10–13]), yields the specific test:

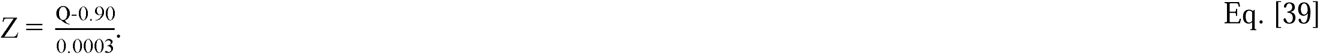

We set 0.05 as the one-sided significance level; thus, the critical value from the standardized normal is Z_c_ = -1.645. Substituting Z_c_ for Z in Eq. [39] and solving for the critical value (Q = Q_c_) yields

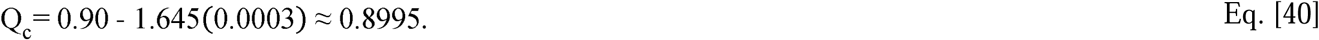

It follows when Q is < 0.8995, the Q-test rejects the null hypothesis multivariate agreement. We note, in X, the observed datasets (bean or mammography) were not multivariate normal by virtue of the univariate marginal pdf comparisons discussed above (i.e., at least one marginal deviated from normality). Nevertheless, the observed data were compared with multivariate normality in both X and Y. As shown in Table 2a, both datasets deviated significantly from multivariate normality in X (Q = 0.07 for the bean dataset and Q ≈ 0.0 for the mammography dataset). In Y, the bean dataset demonstrated normality with Q = 0.97 (i.e., exhibiting a latent normality characteristic), whereas the mammography dataset deviated significantly with Q = 0.83. Multivariate comparison results between the observed and synthetic datasets are shown in Table 2b. The bean dataset was treated with three synthetic models: (A) all components were modeled with KDE; (B) all components were modeled as normal; and (C); first two components were modeled as normal and the third with KDE. For the bean dataset, all three synthetic models in T showed agreement with the observed, whereas models A and C showed considerable but similar disagreement in Y; in contrast, in X (mean-centered), the observed dataset showed multivariate similarity with the respective synthetic datasets for all models. Synthetic mammography datasets showed multivariate agreement with the observed dataset in T, Y, and X. Bean dataset results are also discussed in more detail below. The mammography findings illustrated the approach can reconstruct the multivariate distribution even when a marginal pdf in T with an appreciable variance is non-normal. Thus, the analysis of the mammography data addressed objective one. Likewise, the analysis of the bean data addressed objective two, which is also examined in more detail below.

**Table 2a.** Multivariate normality comparisons in X and Y: the Q test-statistics are provided for the bean and mammography dataset comparisons with multivariate normality. The critical threshold Q_c_ ≈ 0.90.

| Multivariate normality comparison Q test-statistics for observed **x** and **y** |  |  |  |
| --- | --- | --- | --- |
| bean dataset |  | mammography dataset |  |
| **x** | **y** | **x** | **y** |
| 0.07 | 0..97 | $\approx 0.0$ | 0.83 |

**Table 2b.** Observed vs synthetic multivariate comparison Q test-statistics. For the bean dataset: (A) all components were modeled with kernel density estimation (KDE); (B) all components were modeled as normal; (C) first two components were modeled as normal and the third with KDE. For the mammography dataset all components were modeled with KDE. The critical threshold Q_c_ ≈ 0.90.

| Models | Multivariate pdf comparison<br>Q test-statistics |  |  |
| --- | --- | --- | --- |
|  | **t** | **y** | **x** |
| (A) bean dataset | 0.97 | 0.38 | 0.98 |
| (B) bean dataset | 0.97 | 0.98 | 0.99 |
| (C) bean dataset | 0.97 | 0.40 | 0.98 |
| mammography dataset | 0.90 | 0.91 | 0.91 |

### 3.4 Q-test and univariate analysis

The univariate comparison results cited in Table 1b are revisited with a more formal Q-test analysis by treating the outcomes as independent Bernoulli trials. Here, the same one-sided Q-test in Eq. [39] is reformulated for the univariate marginal pdf comparisons by setting p_0_ = 0.90 with N = 1000. These quantities give var(Q) = 9 ×10^-5^, yielding Q_c_ ≈ 0.88. Applying the analogous test to the univariate pdf comparison fractions in Table 1b showed agreement across all components in both the bean and mammography datasets.

### 3.5 Data compression analysis

A more detailed examination of the bean dataset findings in Y and X (Table 2b) is warranted to address objective two. In T, the third component accounted for only a negligible fraction of the variance (approximately 0.01%), which could be considered residue. Thus, the projection test in T was dominated by the first two PCA components and the Q-test showed similarity in all models. Although the residual pdf (3^rd^ variable) showed strong deviation from normality in T, modeling with the non-parametric assumptions (KDE, Model A) led to substantial multivariate disagreement in Y. Similarly, modeling the third component with KDE (non-parametric) while modeling the first two components as normal (Model C), produced comparable disagreement in Y as expected (i.e., KDE replicated the normal pdfs for the first two modes in this Model C, essentially reproducing Model A). Thus, the loss of agreement is driven specifically by how the residual was treated, rather than the dominant modes. In contrast, modeling all components as normal yielded strong agreement with the observed bean dataset in Y and X, as well.

## 4. Discussion and Conclusions

The three methodological objectives produced complementary findings. First, when a non-negligible marginal distribution in T was non-normal, the multivariate distribution could still be synthesized by independently modeling the uncorrelated marginals using KDE or a combination of KDE and normal marginal models. Second, under substantial data compression, residual modes could exhibit non-normal empirical distributions even when the dominant modes were normal; however, synthetic data was better reproduced when these residual modes were modeled as normal, consistent with the residual structure being dominated by compression-related effects; this finding parallels our previous investigation with higher-dimensional data, where modeling the residuals as normal was found empirically to perform well but was not investigated further [30]. Third, the random projection framework was extended from the previously developed multivariate normality application to comparison of two general multivariate distributions [30]. As demonstrated, the resulting two-level testing architecture transforms repeated nonparametric projection-level comparisons into a standardized normal test statistic under a working binomial approximation; this test also provides a means of interrogating differences in multivariate covariance structure as shown theoretically. Although this work was implemented in the low-dimensional regime, the application of optimized univariate KDE at the front end in the X representation is an important feature of the framework. It expands the number of observations available for constructing smooth mappings between X and Y when the number of observations is limited. Taken together, this methodological framework provides an empirical system for evaluating whether the lack of correlation provides an adequate approximation to independence for multivariate reconstruction. It also provides a means of characterizing the residual structure arising under substantial data compression and investigating its appropriate modeling.

The work was motivated in part by the development of a formal testing framework without parametric assumptions for the multivariate distributions under analysis. Our non-parametric test development was based on random projections that reduce a given comparison to scalar quantities. The Q-test is derived from this projection-based framework. Here, multivariate distributional agreement is defined by the frequency of one-dimensional concordance across random directions and multiple replicate comparisons for each direction (i.e., multiple replicate synthetic datasets compared with the observed). The central innovation is the decoupling of the KS test significance level from the final inference, where projection-level KS outcomes are treated as binary evaluations and the Q statistic defined as a functional of their empirical distribution. In this formulation, the KS two-sample test serves only as a local (inner) decision rule, while the global hypothesis test (outer test) is conducted as a normal model approximation for a binomial process independent of the KS test significance level. This methodology produces a scalable test that samples information about the eigen-structure, directional geometry, covariance, and frequency behavior of the multivariate distribution, while not relying on multivariate density estimations or kernel mappings. Using relatively large number of trials provides the basis for invoking a normal approximation to the binomial distribution for the Q-test inference; for reference, we use a conservative lower bound estimate for normality 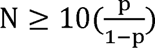 [38] to assess the merits of our applications. Letting p = 0.90, this approximation gives N ≥ 90 as an adequate criterion under the working binomial approximation, not unconditionally; our application far exceeds this bound. Although N was appropriate for the normal approximation, this N does not specify the number of random projection directions required to approximate the Cramér-Wold condition. This projection-based Q-test was applied in X, Y, and T representations allowing multivariate distributional agreement evaluation at each data representation, from the original X representation through marginal normalization in Y to the uncoupled T representation, where the covariance geometry is made explicit. We also showed that the test was invariant with respect to the PCA transformation in theory and practice. Derivations were also provided that showed the impact of applying projections on non-mean center data. Outside of the projection test comparison methodology, the analogous testing framework applies when a sequence of binary outcomes can be modeled as independent Bernoulli trials. Consequently, the same test was reformulated and applied to the univariate marginal comparisons.

We take a slight deviation to explain the bean dataset behavior and the treatment of the residual. Although not explicitly used in this analysis, it is instructive to consider the sequential PCA formulation (i.e., NIPALS approach by Wold [39]) to the same PCA decomposition because it provides a more intuitive interpretation (e.g., see the tutorial in [40]); this approach extracts principal components one at a time and then removing the respective component’s influence from the data matrix making it evident when trailing components reflect negligible residual variation or continued rounding artifacts (this effect is paralleled in Eq. [30]). These artifacts become conspicuous when data shows considerable compression in T, as with the bean dataset. In this context, the final component in T is best interpreted as a near-null direction dominated by numerical residuals arising from data compression, finite precision, and transformation artifacts rather than an informative stochastic mode. Consequently, the apparent non-normality of this component is not structurally informative and is unstable under modeling. These results support the conclusion that in strongly compressed data, residual PCA components should be interpreted as numerical residue and applying flexible nonparametric models, such as KDE, can overfit rounding-level artifacts that degrade multivariate model reconstruction. This interpretation is substantiated by our prior findings, where all residuals were modeled as normal regardless of the observed data characteristics. There was also another inconsistency in the respective comparisons in X for the same bean dataset models. A model that fails comparison in Y but passes in X indicates multivariate agreement is representation-dependent under nonlinear transformations. Marginal pdfs in Y have equal representation, whereas in X they can exhibit wide-ranging behavior in variation and skewness, for example. While the Cramér–Wold theorem guarantees agreement across all projections characterizes a distribution, it does not imply equivalence across non-linear transformations. In this case, the transformation to Y provides a joint structure that exposes discrepancies that are not present or detectable X; we posit, it is not clear if this artifact is material when data modeling is performed in X. Such differences require evaluation in the predictive modeling context to fully understand.

This study was designed to examine the synthetic generation process and the random projection comparison method in low-dimensional datasets where the X, Y, and T representations could be evaluated explicitly. It also examined when uncorrelated PCA components can be modeled independently and how a low-variance residual affects reconstruction. The generator was fitted to the observed datasets used in these comparisons; therefore, the reported Q values describe agreement with those datasets, rather than performance against an independent sample from the underlying population. These datasets (n = 1334 and d = 3) were selected for direct examination of the method; they do not establish its performance when n is small relative to d.

There are several qualifications related to our methods and findings that warrant discussion. The multivariate Q-test was derived under an independence assumption for projection-level outcomes. While this assumption is only approximate due to possible shared data dependence, it does not alter the individual projection-based comparisons, although it could influence the binomial approximation used for the outer Q-test inference. At this juncture, the degree of dependence among the random projection outcomes has not been established empirically; thus, assuming substantial dependence would be no more justified than assuming independence without further investigation. The parameter p is an empirical design factor that defines the target agreement. This quantity requires further evaluation to determine the appropriate level of stringency in the context of using synthetic data in prediction modeling; in this current work, it was set a priori based on heuristics applied previously. Sensitivity may be improved by changing the computational effort toward a larger number of projection directions and fewer replicates per direction. It is worth noting the well-known problem for sufficiently large N, negligible differences will lead to rejection, which could make the test overly stringent without assessing whether differences are interesting [41]. Thus, the appropriate aggregate N is also a design parameter that requires further investigation. Similarly, more powerful univariate tests along each projection may be appropriate, such as the Anderson-Darling (AD) method that puts more weight on the tails of the distribution in comparison with the KS test [34]. Moreover, as an improvement to capture a broader aspect of the univariate comparison within each projection, a test derived from combining p-values from the AD and KS tests with equal weights will be studied in future work.

The work developed a multivariate testing framework, showed the synthetic generator could accommodate non-normal marginal modeling in T, and provided a detailed analysis of the residue when the observed data shows considerable compression. Future work will evaluate this methodology across diverse datasets in the synthetic data modeling context, further investigate the design parameters p, R, and K and their relationship to predictive modeling outcomes, and characterize the degree of dependence among random projection outcomes and its implications for Q-test inference. We have previously investigated randomly selected datasets, including higher-dimensional data, and found that, to good approximation, they exhibited latent multivariate normality, consistent with the bean data in this report. Whether this behavior is typical or incidental remains unclear; quantifying its prevalence across diverse data types is the focus of ongoing work.

## 5. Declarations

### Author contributions

JH, primary and corresponding author, designed the approach, developed code, and directed the experiments. EF is a co-author, assisted in algorithm and investigation designs, performed most of the coding tasks, applied the statistical tests, and developed the graphics in accord with JH, SE and MS. SE and MS are co-authors, both of whom provided the required statistical and mathematical expertise for the investigation.

### Internal Review Board

All methods were carried out in accordance with relevant guidelines and regulations. All procedures were approved by the Institutional Review Board (IRB) of the University of South Florida, Tampa, FL, IRB # 104715_CR000002. Mammography data was collected either (1) retrospectively on a waiver for informed consent, or (2) prospectively by gaining informed consent approved by the IRB of the University of South Florida, Tampa, FL under protocol 104715_CR000002. H. Lee Moffitt Cancer Center & Research Institute (MCC) does not operate its own IRB. However, MCC maintains a Federal Wide Assurance (FWA, 00001464) designating several accredited IRBs to ensure proper oversight of research involving human subjects. The relationship between each IRB and MCC is governed by an affiliation agreement or contract; MCC has such an agreement with the IRB referenced above. Consequently, all research was performed at MCC.

### Use of Artificial Intelligence

Artificial intelligence (AI) was not used to compose this manuscript. AI was used in part to search for relevant research articles that were then inspected by the authors for relevance.

### Data Sharing

Image and clinical data used for this work were made public by JH. Data can be found at this reference [37]. The summarized data and kernel weights used in the analyses are provided as Supplementary Information. Code for this work is publicly available: https://github.com/erinfowler-moffitt/Moffitt-Cancer-Center-Tampa-FL-Heine-Lab/tree/main/projects/automated-breast-density-detection-idl. The algorithms and statistical procedures are described in sufficient detail in this manuscript and the referenced work to support replication. The corresponding author may be contacted for additional information or assistance with the implementation and replication of this work.

